# Widespread occurrence of botulinum and tetanus neurotoxin genes in ancient DNA

**DOI:** 10.1101/2025.02.24.639866

**Authors:** Shyan Mascarenhas, Harold P. Hodgins, Andrew C. Doxey

## Abstract

**Background:** Ancient DNA collected from archaeological specimens not only provides a window into ancient human genetic diversity but also contains a rich mixture of associated microbial DNA including potential pathogens. In recent work, we identified *C. tetani* and tetanus neurotoxin (TeNT) genes in ancient DNA datasets collected from human archaeological specimens. However, the reasons underlying the occurrence of these toxin genes and the extent to which other toxin genes are present in ancient DNA is unclear.

**Methods:** Here, we performed a large-scale analysis of 6,435 ancient DNA (aDNA) sequencing datasets including human and non-human sources, searching for 49 clostridial neurotoxin types and subtypes, as well as 3 additional unrelated toxins.

**Results:** Our search identified a total of 105 ancient DNA datasets (1.6%) containing significant matches to one or more neurotoxin genes. Consistent with our earlier work, TeNT genes were most common, found in 50 ancient DNA datasets. In addition, we identified sequences encoding diverse botulinum neurotoxins including BoNT/C (40 samples), BoNT/D (6 samples), BoNT/B (4 samples), BoNT/E (1 sample), and the *Enterococcus*-associated BoNT/En (10 samples). TeNT genes were detected in a broad range of ancient samples including human and animal (horse, wild bear, chimpanzee) remains, whereas the largest diversity of toxins was detected in aDNA from Egyptian mummies. Phylogenetic and sequence analysis of the identified matches revealed close identity to modern forms of these toxins. Damage analysis revealed several toxin genes with hallmarks of ancient DNA associated damage, indicative of an ancient origin.

**Conclusions:** Our work reveals that clostridial neurotoxin genes occur frequently in aDNA samples, including human and animal-associated toxin variants. We conclude that the frequent association of these genes with aDNA likely reflects a strong ecological association of pathogenic clostridia with decaying human and animal remains and possible post-mortem colonization of these samples.

## Introduction

Clostridial neurotoxins (CNTs), which include botulinum neurotoxins (BoNTs) and tetanus neurotoxins (TeNTs) are the most potent toxins known to science, and causative agents of botulism and tetanus neuroparalytic diseases (1). BoNTs and TeNTs, typically produced by *Clostridium botulinum* and *Clostridium tetani*, respectively, are highly potent bacterial neurotoxins that interfere with synaptic transmission by cleaving key proteins in the SNARE complex, essential for neurotransmitter release (2, 3). Both toxins are zinc-dependent metalloproteases that specifically target different SNARE proteins, affecting distinct parts of the nervous system and producing opposite physiological outcomes (4–6).

BoNT consists of a heavy and light chain linked by a disulfide bond. Upon entry into the peripheral motor neurons via receptor-mediated endocytosis, the heavy chain facilitates translocation of the light chain into the cytoplasm (7, 8). The light chain then cleaves SNARE proteins, such as SNAP- 25, VAMP/synaptobrevin, or syntaxin, depending on the BoNT serotype, preventing the release of acetylcholine at the neuromuscular junction. This inhibition of acetylcholine release leads to flaccid paralysis, as the muscle fibers cannot contract without stimulation.

In contrast, TeNT follows a similar route of entry but exhibits a different cellular trafficking pattern (6). TeNT is taken up by peripheral neurons and undergoes retrograde transport to the spinal cord and brainstem, where it specifically targets inhibitory interneurons (9, 10). Once inside these neurons, TeNT’s light chain cleaves VAMP/synaptobrevin in the SNARE complex, preventing the release of inhibitory neurotransmitters such as glycine and GABA (11). This blockage of inhibitory signaling leads to continuous excitatory stimulation of motor neurons, causing the characteristic spastic paralysis observed in tetanus, with muscle rigidity and spasms.

### The growing diversity of BoNT types and BoNT-related sequences

BoNTs exhibit remarkable diversity, with multiple distinct types (BoNT/A to BoNT/X) identified based on sequence variations, receptor specificity, and differences in protease activity (12–16). Each BoNT type, produced by various strains of *Clostridium botulinum* and related *Clostridium* species, possesses unique functional properties that enable binding and entry into specific neuronal subtypes, contributing to variations in potency and clinical application. The well-characterized BoNT serotypes A, B, E, and F are associated with human botulism, while others, such as BoNT/C, D, and G, affect animals or have less well-defined roles in human disease. BoNT/X, discovered more recently through bioinformatics analysis, cleaves a distinct set of SNARE proteins, including SNAP-47, broadening our understanding of BoNT target diversity and cellular impact.

Recent advances in bioinformatics and genomic sequencing have led to the identification of a wide range of BoNT-related proteins (BoNT-like proteins) in diverse bacterial genera, often through homology searches and domain-based analysis (14, 17–25). These BoNT-like proteins contain similar structural features, such as the heavy and light chain architecture and zinc metalloprotease activity, yet they exhibit variations in substrate specificity and cellular effects. For example, homologs identified in non-clostridial species such as *Weissella* (19, 26) and *Enterococcus* (22, 23) possess BoNT-like domains and protease activity but their host specificity and neurotoxicity remains unclear. The recent identification and characterization of PMP1 (24), a BoNT-related neurotoxin that targets Anopheles mosquitoes, highlights the potential association of these more divergent BoNT-like toxins with invertebrate hosts.

### Extreme rarity of CNTs in sequencing datasets and unexpected identification in ancient DNA

Building on the bioinformatic above, researchers have also performed large-scale systematic searches of diverse metagenomic datasets, expecting to find CNT genes frequently in environments like soil, food, and animal-associated microbiomes, where clostridia are commonly found (17). Surprisingly, however, such analyses have revealed that CNT genes are virtually undetectable in available environmental metagenomic datasets (including soil), despite the ubiquity of clostridia in these environments.

Recently, Hodgins et al., performed an “unbiased” search of the entire NCBI Sequence Read Archive for *C. tetani*, with the goal of identifying datasets and potential new niches harboring TeNT genes (27). Unexpectedly, this study led to the identification of *C. tetani* genomes (and TeNT genes) in 38 ancient DNA datasets collected from a wide array of different archaeological specimens associated with different geographical regions and time points (27). This study raises several intriguing questions: 1) How prevalent is toxigenic *C. tetani* in ancient DNA collected from archaeological samples? 2) What is the underlying reason for the association between *C. tetani* and archaeological DNA? 3) Are additional clostridial pathogens and neurotoxins also prevalent in these datasets? 4) Are these identified sequences indicative of ancient pathogens that were contemporaneous with these historical samples or do they reflect later post-mortem colonization?

To investigate these questions, in this work we perform a large-scale survey of all clostridial neurotoxin genes across thousands of ancient DNA datasets. Our results reveal an even wider diversity of clostridial neurotoxin genes in these samples, and provide some insights into the above questions.

## Methods

### Overview

To identify toxin genes in ancient DNA (aDNA) samples, we constructed a general analysis pipeline shown in **Figure 1**. Our pipeline began with the construction of a “query dataset” of toxin genes as well as a “target database” of aDNA sequencing datasets derived from the NCBI sequence read archive (SRA) (28) to search against. All SRA datasets from the target database were downloaded, processed, and searched by mapping raw reads to reference gene sequences in the query dataset. From the read-pileups, we then computed the consensus sequence of mapped reads to reconstruct the toxin gene sequence for a given SRA dataset. All aDNA-derived toxin sequences were then compared to their closest reference sequences in the query dataset to calculate their percentage identities and coverage (fraction of reference sequence covered by the alignment). In addition, a subset of aDNA-derived toxin sequences was further analyzed by mapDamage (29) to quantify the damage level found in the associated reads, high levels of which are indicative of genuine ancient DNA. The details of this pipeline are described below.

**Figure 1.**
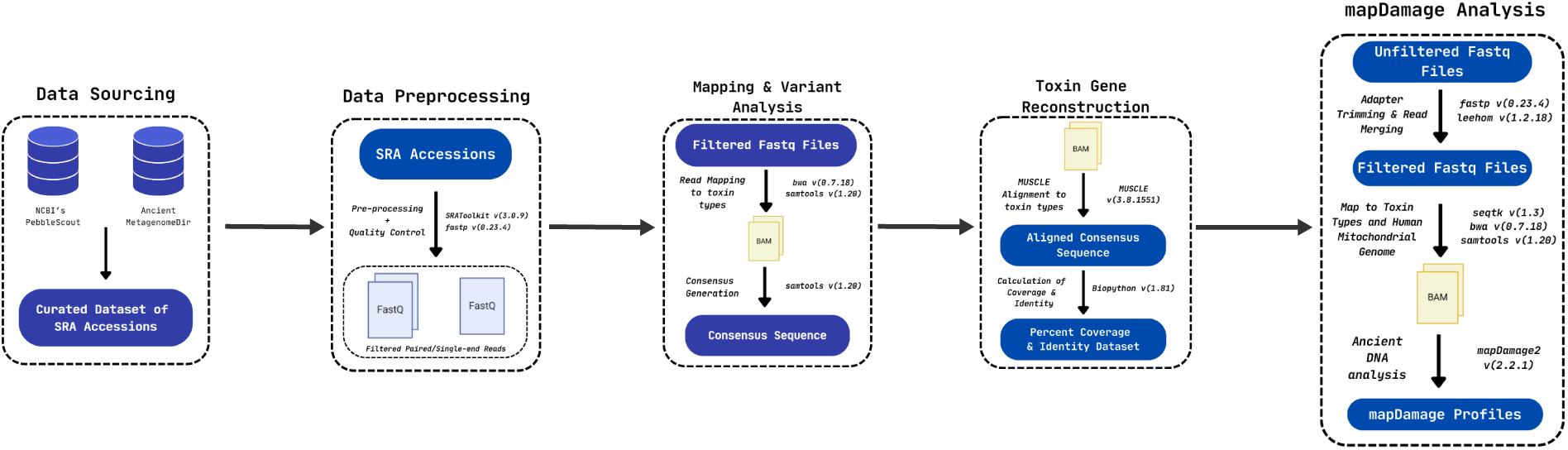
Bioinformatic workflow for identifying toxin genes in ancient DNA (aDNA) datasets from the SRA and ENA. A curated set of aDNA datasets was generated from two sources, and all datasets were screened for the presence of reads mapping to a set of query toxin genes consisting of all clostridial neurotoxin types and three additional toxins for comparison. Consensus sequences of read pileups for toxin-positive datasets were used to reconstruct aDNA-derived toxin sequences. These sequences were further analyzed by aligning them to their closest reference sequence and calculating coverage and percent identity. Finally, mapDamage analysis was performed on a subset of toxin-positive datasets with coverage exceeding 20%.

### Construction of query toxin dataset

We constructed a query dataset of gene sequences for all known CNT types and subtypes (15) including BoNT/A (8 subtypes), BoNT/B (8 subtypes), BoNT/C, BoNT/CD Mosaic, BoNT/DC Mosaic, BoNT/D, BoNT/E (12 subtypes), BoNT/En from *Enterococcus* (23), BoNT/F (9 subtypes), BoNT/G, BoNT/HA/FA, BTNT from *Bacillus toyonensis* (21), BoNT/PMP1 from *Paraclostridium bifermentans* (24), BoNT/TeNT, BoNT/Wo from *Weissella oryzae* (19), and BoNT/X (30) toxins. These sequences were derived from https://bontbase/org. In addition, we also included three query toxins unrelated to BoNT (diphtheria toxin, *C. difficile* TcdB, and anthrax lethal factor), due to ongoing research interests in exploring the evolutionary diversity of these toxins (31–34). See **Supplementary Table 1** for a list of all query toxin sequences.

### Screening of ancient DNA datasets for initial toxin hits

To construct a target database of ancient DNA sequencing datasets for querying, we combined two data sources. First, we sourced datasets from the community-curated AncientMetagenomeDir database (35). The accession numbers were organized into four collections: Ancient Single Genome - Host Associated, Ancient Metagenome - Host Associated, and Ancient Metagenome - Environmental. In total, these collections consisted of 6,404 datasets: 2,931 “Ancient Single Genome - Host Associated” datasets, 2,651 “Ancient Metagenome - Host Associated” datasets, 822 “Ancient Metagenome – Environmental” datasets. For all AncientMetagenomeDir SRA/ENA datasets, we used the Logan (36) tool to pre-screen the datasets for contigs containing matches to any of the query toxin genes. Contigs were aligned to the query toxin gene sequences using Samtools v1.13 (37) and Minimap2 v2.24-r1122 (38). Only those SRA/ENA datasets with contig matches to query toxins were analyzed further.

As a second data source, we used the NCBI’s PebbleScout tool (39) to identify datasets in the NCBI SRA containing matches to *k*-mers found in the toxin queries, performing a separate PebbleScout search for each query. PebbleScout search parameters were set to default, with a masking threshold of 5,663, and a score constant of 13,000. The PebbleScout search identified an additional 31 datasets containing *k*-mers matching the toxin queries (>=1% coverage), bringing the total number of aDNA datasets analyzed in this project to 6,435. All datasets searched are listed in **Supplementary Table 2**.

### Analysis of toxin variants in aDNA datasets

A total of 125 datasets with “initial hits” to toxin genes were further analyzed to gain a more precise understanding of toxin composition and subtypes present in these samples. The SRA-toolkit v3.0.9 was used to download and split SRA accessions accordingly into paired or single-end reads. The reads were then pre-processed using fastp v0.23.4 (40), with default settings, to perform initial quality filtering and remove potential adapter sequences. Following pre-processing, the reads were mapped to gene sequences of all toxins in our query dataset using BWA v0.7.18 and Samtools v1.20. Using samtools v1.20, we then examined the consensus sequence of the read pileups and calculated coverage for each reference toxin. To determine percent identity, we aligned the consensus sequences to their respective reference toxins using MUSCLE v3.8.1551, then calculated their percent identity with Biopython v1.81. In cases where there were multiple toxin types detected, we performed a manual analysis to distinguish genuine “mixed” samples from overlapping predictions due to sequence similarities between closely related types (e.g., BoNT/C and BoNT/D).

### Damage analysis

For the identified toxin genes and datasets identified above, ancient DNA damage analysis was performed using mapDamage (29). The original FASTQ files were first processed using leehom v1.2.18 (41) and fastp v0.23.4 (40). Short sequences under 30 bp were filtered out using seqtk v1.3 with the command *seqtk seq -L30*. Because leeHom v1.2.18 merges paired-end reads, datasets processed with leeHom were mapped as single-end reads to the identified toxin type(s) using BWA v0.7.18 with the *bwa samse* command, as done previously (27). In contrast, datasets filtered with fastp v0.23.4 retained paired-end reads, which were mapped to the identified toxin type(s) using BWA v0.7.18 with the *bwa sampe* command, also following (27). For comparison, reads were also mapped to the human mitochondrial reference genome (accession: NC_012920.1). BAM files were sorted with samtools v1.20, and misincorporation rates were calculated using mapDamage v2.2.1 with options --merge-reference-sequences --no-stats.

### BLAST analysis of reconstructed toxin genes from aDNA

All aDNA-derived toxin gene sequences were then aligned to the NCBI-nr database, which allowed us to both verify their identity and compare them to the full diversity of toxin sequences in Genbank. Because most aDNA-derived toxin sequences were fragmented and contained gaps, we split them into contiguous fragments and BLASTed each fragment individually. All identified SNPs were recorded and numbered using a common reference gene for each toxin type. A lollipop SNP plot was then generated using ggplot2.

## Results

### Ancient DNA datasets harbor diverse types of clostridial neurotoxins

Using the bioinformatics tools pebbleScout (39) and Logan (36), we searched a total of 6,435 ancient DNA (aDNA) sequencing datasets for sequences matching known clostridial neurotoxin types as well as three unrelated toxins - diphtheria toxin, anthrax lethal factor, and the *C. difficile* toxin, TcdB. Our search identified 125 “candidate” datasets containing potential matches to query toxins. We then reconstructed toxin sequences from these aDNA datasets by mapping reads to a non-redundant set of reference toxin genes (see **Figure 1**). A total of 105 aDNA datasets (1.6% of the original 6,435) had reads mapping to one or more query toxin gene, each of which was further validated by BLAST analysis (**Figure 2A**). In total, TeNT sequences were identified in 50 ancient samples, BoNT/C in 40 samples, BoNT/En in 10 samples, BoNT/D in 6 samples, BoNT/B in 4 samples, and BoNT/E in 1 sample (**Figure 2A**). We also detected a mixture of more than one toxin type in 19 samples (21%) (**Figure 2B**). For example, we detected both TeNT and BoNT/D (at lower relative abundance) in sample “Abusir1655-tooth”, as can be seen visually in read pileups (**Supplementary Figure 1**). No matches to the other BoNT types or three non-BoNT toxins were detected.

**Figure 2.**
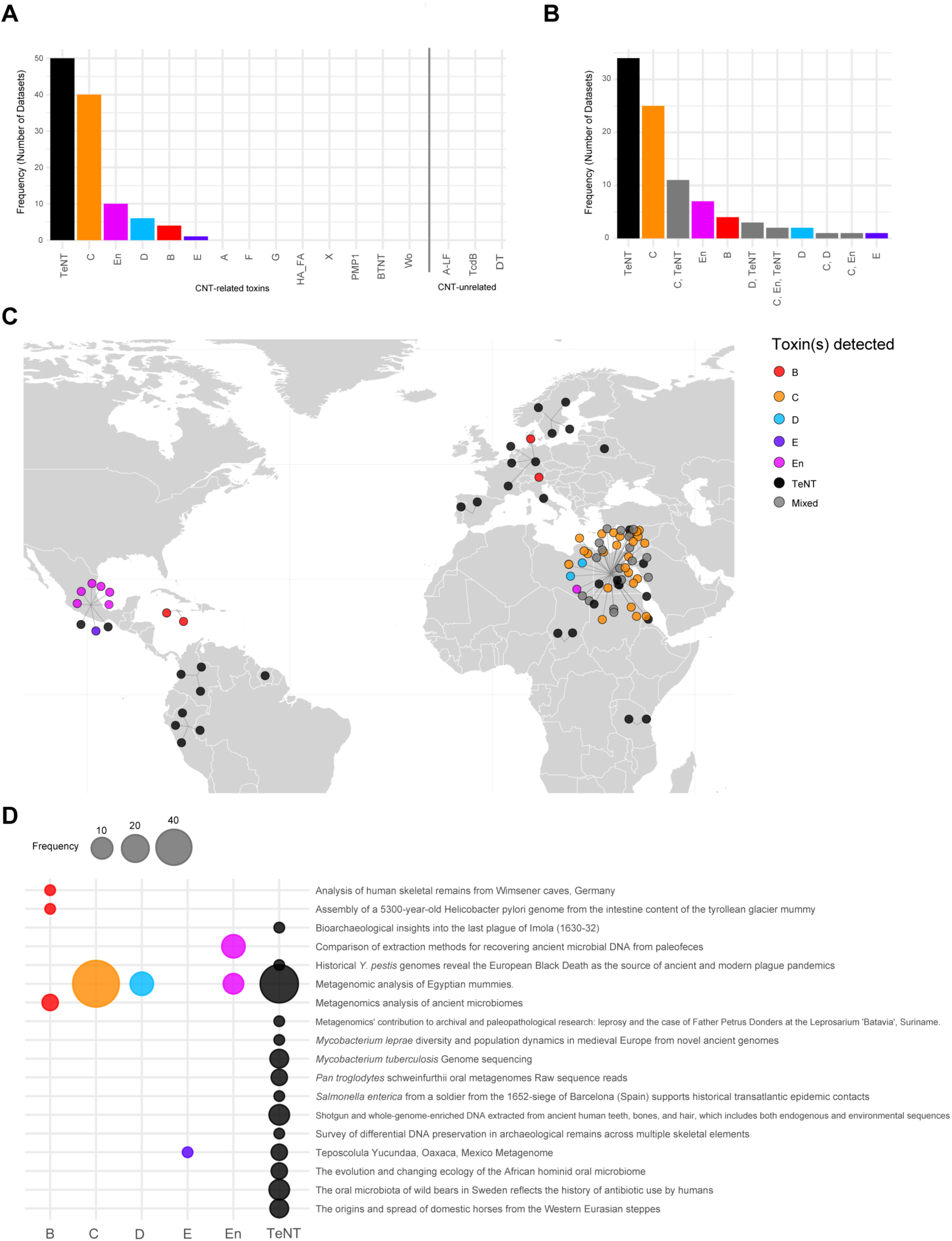
Diversity and distribution of neurotoxin genes detected in ancient DNA datasets. A) Frequency of toxin genes detected across different ancient DNA datasets (BioSamples). B) Number of aDNA datasets identified containing different toxin gene combinations. C) Geographic distribution of toxin-positive aDNA datasets. D) Frequency of toxin genes detected across unique ancient DNA studies.

### Clostridial neurotoxin genes are present in ancient DNA associated with different host species and geographic regions

The 105 aDNA datasets with matches to clostridial neurotoxin genes (“toxin positive” datasets) are derived from 91 distinct ancient samples from geographically diverse regions spanning four contents (**Figure 2C**) and 18 separate studies (**Figure 2D**). As shown in **Figure 2C** and **2D**, the largest quantity (n = 54) and diversity of toxin genes (6 distinct types) were detected in aDNA from Egyptian mummy samples (42). Among all the toxin types, TeNT genes were most broadly detected across the studies (14 of 18). Interestingly, TeNT genes were detected not only in aDNA associated with human archaeological remains, but also aDNA from domestic horses (43), oral microbiota from wild bears (44) and chimpanzees (45). Additional metadata associated with toxin-positive datasets is included in **Supplementary Table 3**.

### Clostridial neurotoxin genes from ancient DNA datasets are highly similar to modern reference sequences

Next, we examined the degree of sequence similarity between the identified neurotoxin sequences in aDNA and their respective reference genes. The neurotoxin genes detected across the 91 aDNA samples are displayed in **Figure 3A**, with their coverage (**Figure 3B, Supplementary Table 4**) and percentage identities (**Figure 3C, Supplementary Table 5**). These toxin genes are highly similar to the reference genes, ranging from 96% to 100% sequence identity. However, it is important to note that the toxin genes associated with lower percentage identities below 98% were partial sequence fragments with alignment coverage below 10% (**Supplementary Figure 2**), which may result in poor estimates of their true similarity. These partial sequences were predominantly associated with BoNT/B, BoNT/E, and BoNT/En (**Figure 3B**). However, higher coverage matches were obtained for BoNT/C (max = 97.9%), BoNT/D (max = 47.1%) and TeNT (max = 99.5%) (**Figure 3C**). Again, all sequence matches to BoNT genes (including fragments) were validated through subsequent BLASTn analysis of the full ncbi-nr database, which confirmed their identity as BoNT sequences. An example read pileup is shown in **Figure 3D**, which depicts a high-coverage detection of BoNT/C in an Egyptian mummy ancient dataset (SAMEA6847470).

**Figure 3.**
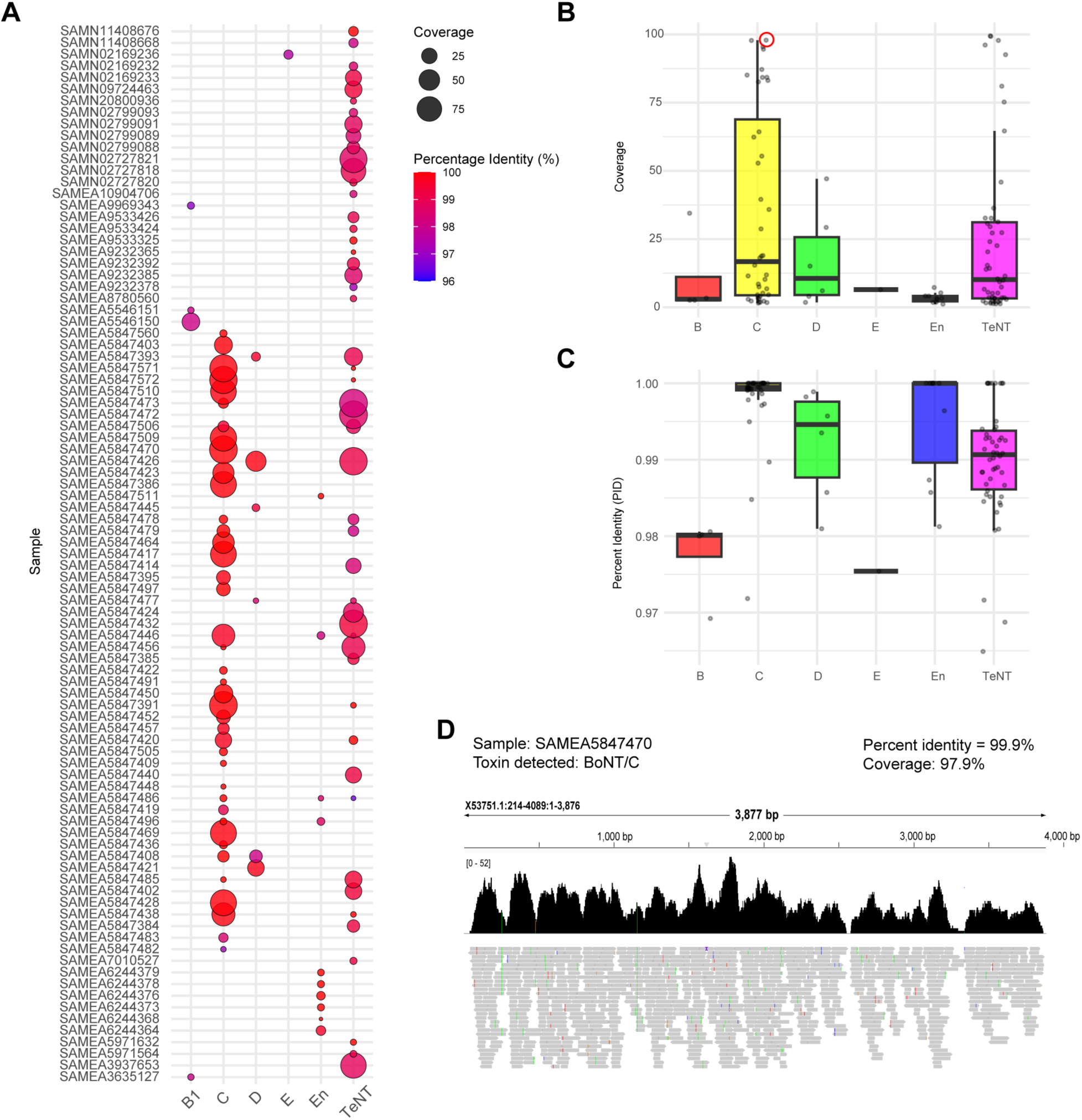
Analysis of neurotoxin genes detected in 105 ancient DNA datasets. A) Bubble plot visualization of neurotoxin genes detected in 105 aDNA datasets, along with their percentage identity and coverage relative to modern reference sequences. Their coverage distributions (B) and percentage identity distributions (C) are also shown as boxplots per toxin. D) A read pileup for an example high-coverage dataset containing BoNT/C reads (circled in B). The pileup shows the individual reads mapped to the BoNT/C reference gene sequence and read density. As shown, there is broad coverage of BoNT/C in this dataset with reads mapping across the entirety of the BoNT/C gene. See **Supplementary Table 3** for accession #s of the 105 datasets listed in (B).

### Ancient DNA datasets contain novel variants of clostridial neurotoxin sequences

Despite being highly similar to modern reference genes, it is possible that the aDNA-derived neurotoxin genes possess unique sequence variants. To investigate this further, we performed a BLAST comparison of all aDNA-derived toxin sequences to the full diversity of neurotoxin sequences available in NCBI/GenBank, excluding TeNT sequences as this has been investigated previously (27). The analysis revealed a total of 28 aDNA-derived toxin genes that possess 47 SNPs not found in modern toxin sequences (**Table 1**, **Figure 4**). The most frequent SNPs detected were G→A (n = 8), C→T (n = 8), A→G (n = 8), T→C (n = 7), and C→A (n = 7) mutations. Importantly, some of the identified SNPs may be sequencing errors due to ancient DNA associated damage; particularly G→A and C→T. Visualization of these SNPs across the full-length sequences reveals that variants occurred sporadically throughout identified BoNT/B, BoNT/En, and BoNT/C toxins (**Figure 4**). However, identified mutations in BoNT/D variants were concentrated in the middle region of the sequence and 3’ region. The largest number of observed unique mutations occurred in a BoNT/D sequence from ERR4374018 (n = 10 SNPs).

**Figure 4.**
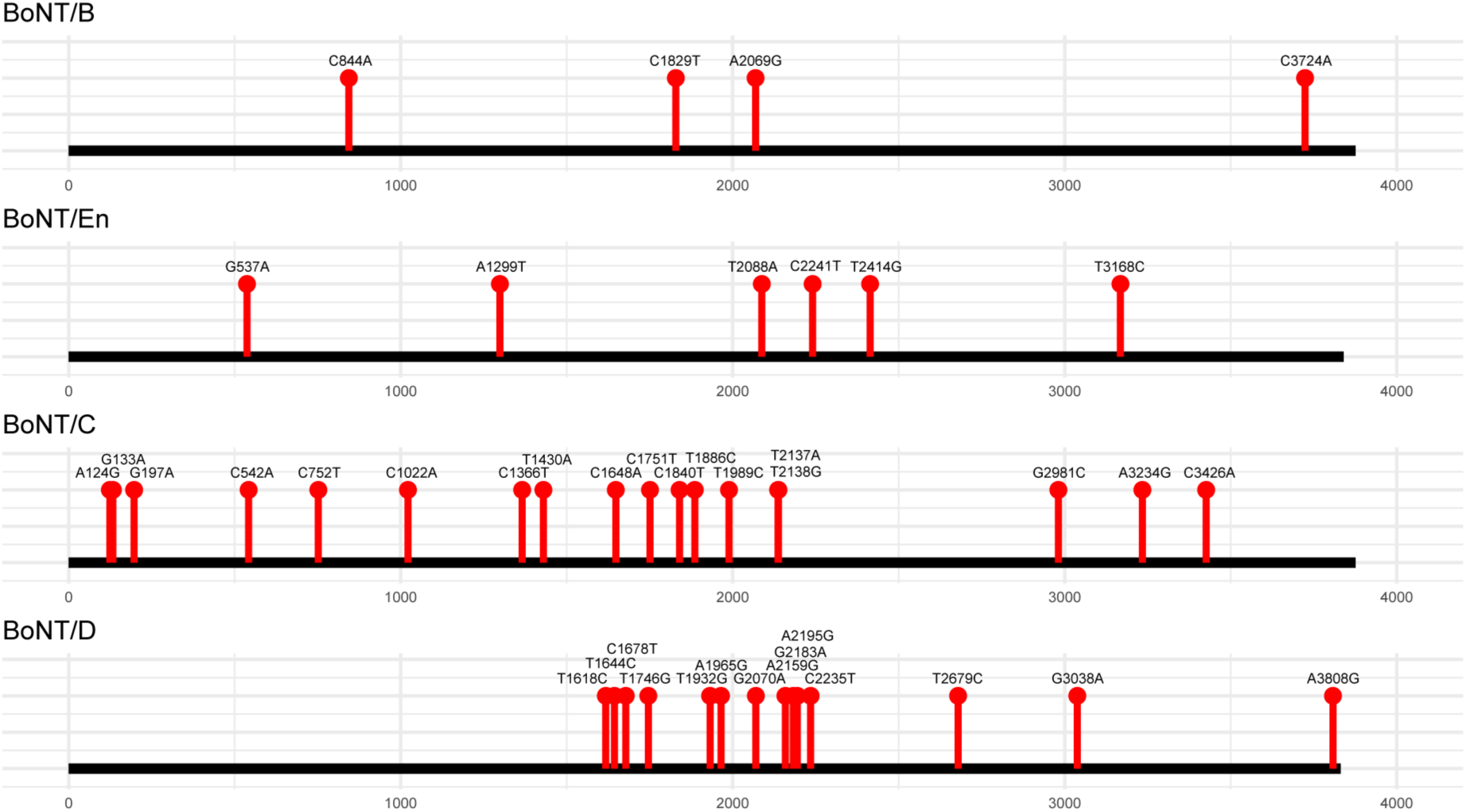
Putative neurotoxin SNPs identified in ancient DNA datasets. Some SNPs detected may be the result of DNA damage associated mutations. TeNT sequences were not included as a thorough analysis has been performed elsewhere (27).

**Table 1.**
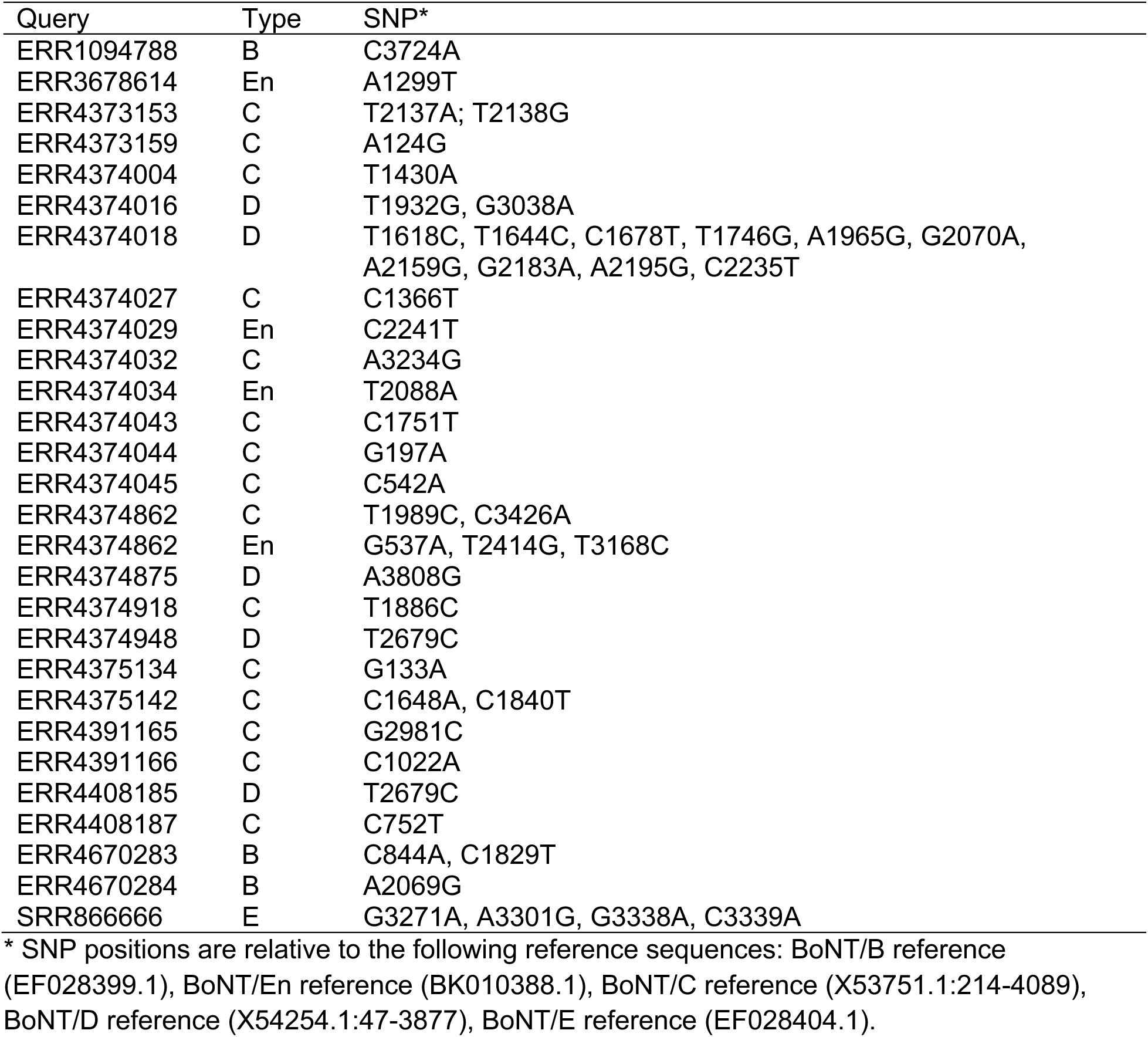
Putative SNPs in neurotoxin genes detected in ancient DNA datasets.

**Table 2.**
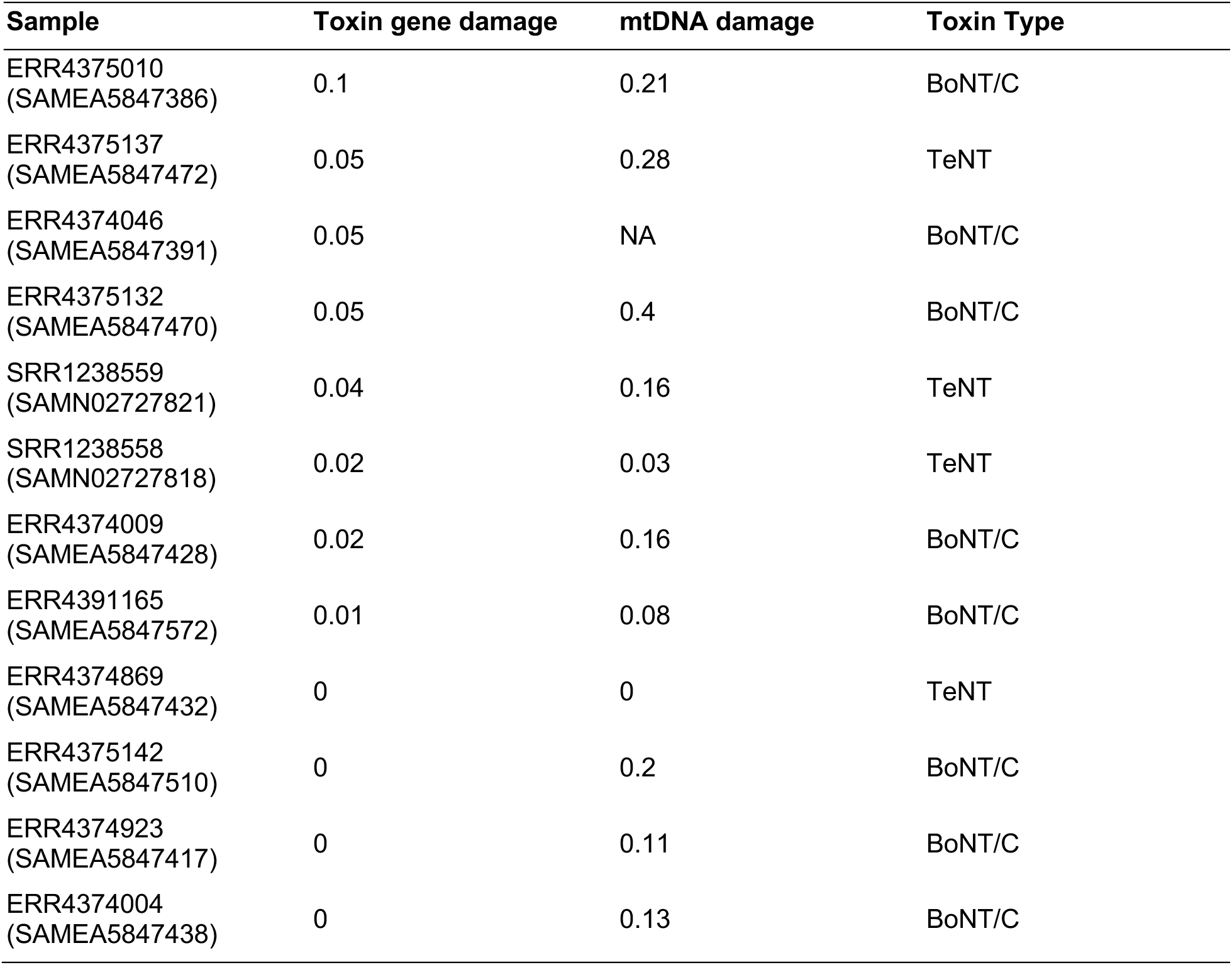
Damage levels associated with toxin genes identified in ancient DNA datasets.

| Sample | Toxin gene damage | mtDNA damage | Toxin Type |
| --- | --- | --- | --- |
| ERR4375010<br>(SAMEA5847386) | 0.1 | 0.21 | BoNT/C |
| ERR4375137<br>(SAMEA5847472) | 0.05 | 0.28 | TeNT |
| ERR4374046<br>(SAMEA5847391) | 0.05 | NA | BoNT/C |
| ERR4375132<br>(SAMEA5847470) | 0.05 | 0.4 | BoNT/C |
| SRR1238559<br>(SAMN02727821) | 0.04 | 0.16 | TeNT |
| SRR1238558<br>(SAMN02727818) | 0.02 | 0.03 | TeNT |
| ERR4374009<br>(SAMEA5847428) | 0.02 | 0.16 | BoNT/C |
| ERR4391165<br>(SAMEA5847572) | 0.01 | 0.08 | BoNT/C |
| ERR4374869<br>(SAMEA5847432) | 0 | 0 | TeNT |
| ERR4375142<br>(SAMEA5847510) | 0 | 0.2 | BoNT/C |
| ERR4374923<br>(SAMEA5847417) | 0 | 0.11 | BoNT/C |
| ERR4374004<br>(SAMEA5847438) | 0 | 0.13 | BoNT/C |

### Damage analysis of toxin genes reconstructed from ancient DNA

Characteristic patterns of post-mortem DNA damage include increased cytosine deamination near fragment ends, which results in C→T (or G→A) substitutions (46). To investigate whether aDNA-derived toxin sequences display evidence of ancient DNA damage, we used mapDamage (29) to quantify the misincorporation rates in reads. For comparison, we also analyzed the damage levels of human mitochondrial DNA (mtDNA) from the same samples.

Only 12 of the identified aDNA-derived toxin genes had sufficient depth of coverage for analysis by mapDamage. As shown in Table 1, human mtDNA in 7 of these 12 (58%) samples had significant damage (5’ misincorporation rate exceeding 10%). However, only one of the identified toxin genes (*bont/c* in sample SAMEA5847386, **Figure 5**) had a significant damage level of 10%. Four of the toxin genes had no damage detected, and seven had lower damage levels ranging from ∼1% to ∼5%. We further investigated the putatively ancient *bont/c* gene detected in SAMEA5847386 using an alternate aDNA workflow (see Methods). This analysis revealed a damage rate exceeding 10% and a shifted read length distribution indicative of genuine ancient DNA (**Supplementary Figure 3**).

**Figure 5.**
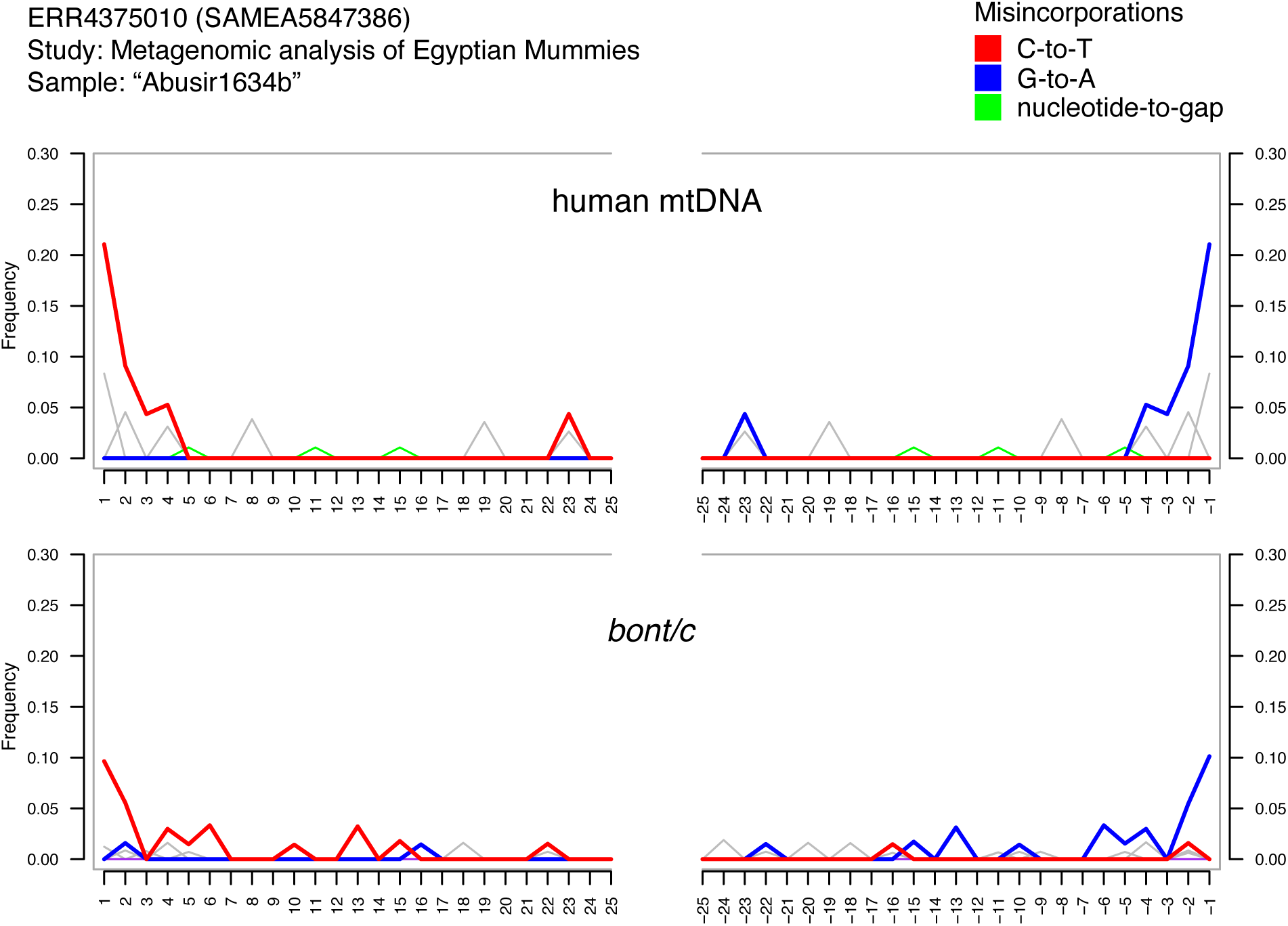
MapDamage profile for reads matching *bont/c* (bottom) and mitochondrial DNA (top) from an Egyptian mummy aDNA dataset. Reads were first processed using fastp v0.23.4. The increase in C→T (and G→A) misincorporations at the edges of reads is a hallmark of ancient DNA. The first position damage level of 21% (mtDNA) and bont/c (10%) is above previous thresholds established for ancient DNA authentication.

## DISCUSSION

In previous work, we identified and reconstructed *C. tetani* genomes and associated *tent* genes in aDNA datasets collected from 38 diverse human archaeological samples (27). These findings raised several questions (as described in the introduction) such as whether ancient DNA samples harbor additional types of toxin genes, and the underlying reasons for the occurrence of neurotoxins in archaeological DNA. To investigate these questions, here we investigated the presence of a variety of toxin genes including all clostridial neurotoxin types in 6,435 aDNA datasets. Our survey identified neurotoxin sequences in 1.6% (105 datasets) of the total examined, including a broad diversity and geographic range of sample types. Our analysis also revealed a diverse array of neurotoxins, including not only tetanus neurotoxin (TeNT) but also multiple botulinum neurotoxin types such as BoNT/B, C, D, E, and En. Our work therefore suggests that, while uncommon, clostridial neurotoxin genes in general appear to be broadly distributed and statistically enriched in ancient DNA. Our work here is also notable by providing the first identification of BoNT genes in ancient DNA, adding to our previous study on TeNT (27).

The findings reinforce our previous work demonstrating the widespread occurrence of TeNT genes in ancient DNA datasets. TeNT was detected in 50 ancient samples, making it the most prevalent toxin gene identified. Notably, TeNT was not restricted to human-associated archaeological remains but was also identified in aDNA from domesticated horses, wild bears, and chimpanzees. This widespread detection likely reflects the ubiquity of *C. tetani* across diverse hosts and environments (47). The frequent detection of TeNT in multiple ancient host-associated contexts potentially reflects a historical role in animal and human health, although its presence alone does not indicate disease occurrence.

Unlike TeNT, BoNT genes exhibited more restricted host and geographic distributions. For example, BoNT/B was found only in ancient human samples and not in animals. This restricted occurrence aligns with modern epidemiology, where BoNT/B is associated with human botulism along with A and E (48), rather than environmental reservoirs. Although BoNT/A was not detected, a single case of BoNT/E was detected in an ancient DNA sample from a 16th-century Mixtec individual from Teposcolula Yucundaa (Oaxaca, Mexico), associated with the “huey cocoliztli” unknown disease outbreak (49). The human-specific detection of both BoNT/B and BoNT/E is intriguing given the association between these serotypes and human botulism outbreaks.

Interestingly, BoNT/C and BoNT/D however were found exclusively in Egyptian mummy samples, despite the fact that modern BoNT/C and D are primarily associated with animals, particularly wild birds as well as cattle (50). This unexpected finding raises questions about its ecological and historical significance. It is unclear whether BoNT/C and D were introduced through environmental contamination, were naturally present in ancient Egyptian human microbiomes, or played a previously unrecognized role in human-associated environments.

Fragments of BoNT/En, which was initially identified in an *Enterococcus faecium* strain isolated from cow feces (22, 23), were detected in ancient human paleofeces, marking the first identification of En-related sequences outside of its original discovery. This provides important new evidence that BoNT/En may have had a broader historical distribution than what is currently recognized. Its presence in paleofeces suggests that it was ingested by ancient populations, potentially through foodborne exposure or interactions with animal microbiomes. This finding warrants further investigation into the origins, distribution, and possible historical impact of BoNT/En in ancient human and animal populations.

A key question arising from our findings is whether these toxin genes represent truly ancient genetic material or are the result of modern contamination. Our sequence similarity analysis revealed that most aDNA-derived toxin genes exhibited high sequence identity (96-100%) to modern reference toxin genes. Despite their close similarity, we identified 28 aDNA-derived toxin gene sequences with unique SNPs not found in modern/reference sequences. While some of these mutations could be attributed to post-mortem DNA damage (e.g., the frequent G→A and C→T transitions), the presence of novel sequence variants raises the intriguing possibility that these represent unique lineages of neurotoxin-producing species. These findings warrant further investigation, particularly through deeper sequencing and phylogenetic analyses to determine whether these represent previously unknown toxin-producing strains. It is thus difficult, based on sequence comparison alone, to determine the antiquity of the identified neurotoxin sequences.

As another strategy to assess the authenticity of these sequences as ancient DNA, we examined post-mortem DNA damage patterns and fragment length distributions using mapDamage analysis. Our results revealed that human mitochondrial DNA (mtDNA) exhibited significant damage (>10% misincorporation rates) in 58% of analyzed samples, consistent with expectations for degraded ancient DNA. However, the majority of toxin genes displayed minimal damage, with only one (BoNT/C in sample SAMEA5847386) exceeding the 10% threshold for significant ancient DNA-associated damage. This discrepancy suggests that most toxin genes in our dataset have undergone less degradation than endogenous host DNA. This may be due to differential DNA preservation (*Clostridium* spores may be more resistant to environmental degradation, leading to better long-term DNA preservation). Or this may be due to potential clostridial contamination from nearby soil, human handling, or scavengers, as discussed earlier (27).

There are also several technical limitations of our study that are worth mentioning. First, the large number of aDNA samples that we search required some bioinformatic approaches that favor speed over sensitivity, which may have led to false negative predictions. For example, by searching with Logan’s pre-assembled contigs from SRA datasets, this enabled straightforward and rapid homology search across thousands of datasets. However, as it is well known that only portions of metagenomic datasets can be properly assembled (51), it is possible that sequencing reads associated with toxin genes are present in the unassembled fractions of some datasets, leading to them being missed. Given this technical limitation and also the difficulty in capturing low abundance species and genes in metagenomes (52), our estimate of neurotoxin prevalence across aDNA datasets is likely an underestimate.

Despite these limitations, our study provides compelling evidence that tetanus and botulinum neurotoxin genes are detectable in hundreds of ancient DNA datasets. The distinct host and geographic distributions of BoNT/B, BoNT/C, and BoNT/En raise important questions about their ecological and evolutionary histories. The detection of BoNT/C exclusively in Egyptian mummy samples and BoNT/En in paleofeces highlights the potential for ancient dietary or environmental exposures to neurotoxin-producing *Clostridium* species. Future studies could enhance these insights by exploring deeper sequencing efforts to improve coverage, or metagenomic analysis of ancient fecal and soil samples to better understand the past environmental reservoirs and transmission routes of these neurotoxins. In addition, our bioinformatic workflow can be applied to study the explore the diversity and distribution of other virulence factors or genes of interest in ancient samples.

## Acknowledgements and Funding

This work was supported through an NSERC Discovery Grant as well as a University Research Chair (A.C.D.).

## Author Contributions

ACD was responsible for conceptualization, funding acquisition, project administration, and supervision. ACD and SM were responsible for formal analysis, investigation, and methodology. All authors were responsible for data curation, visualization, and writing.

## Data Availability

All datasets analyzed in this study are publicly available and listed in the Supplementary Material. Source code associated with this project is available in github at: https://github.com/doxeylab/Mascarenhas-aDNA-toxin-survey

---

**Supplementary Figure 1.**
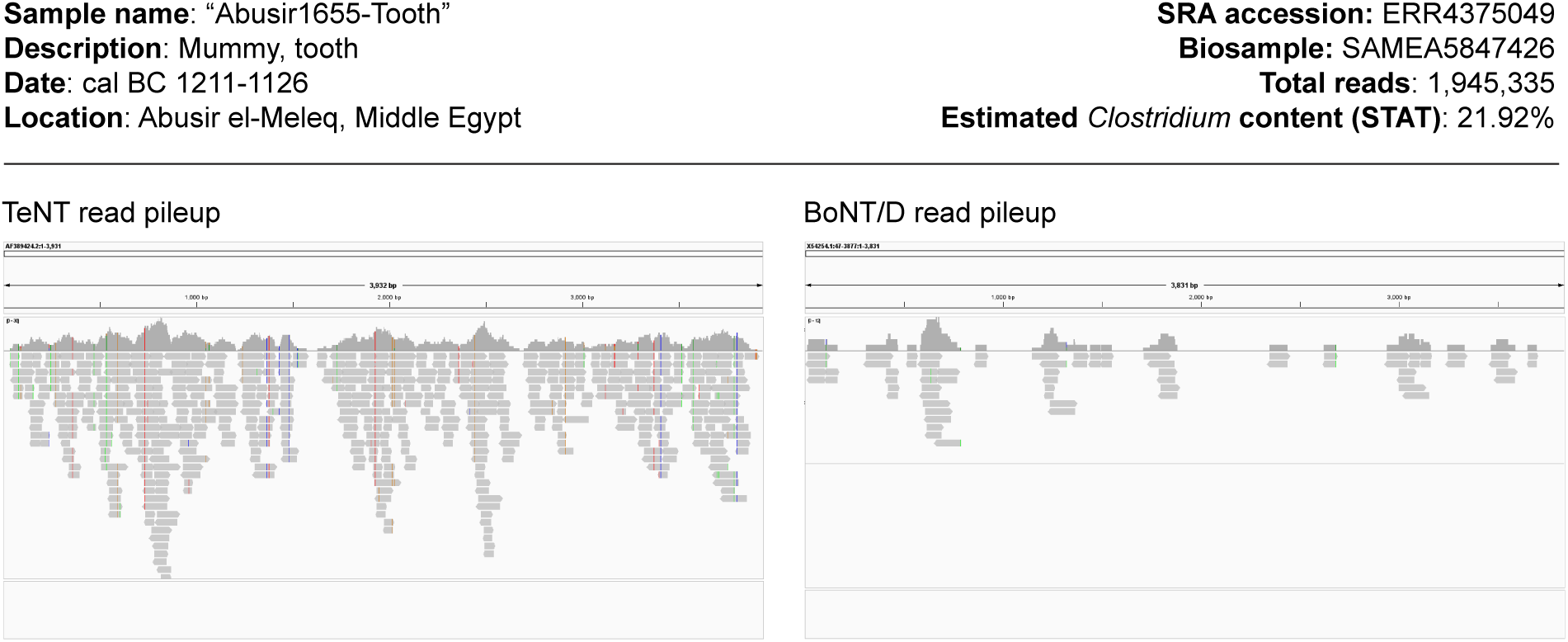
An ancient DNA dataset with reads matching both TeNT and BoNT/D.

**Supplementary Figure 2.**
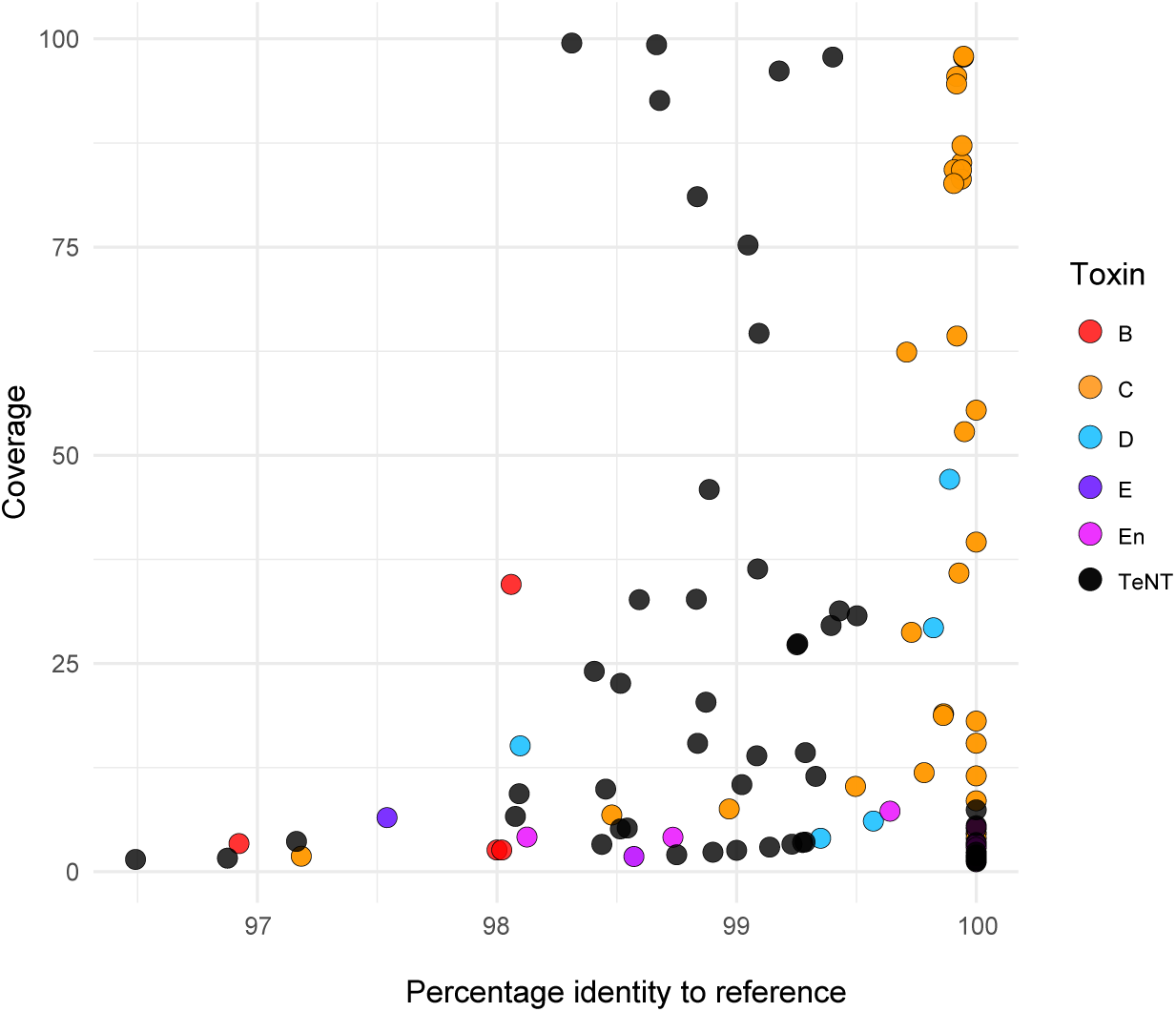
Scatterplot depicting the coverage and percentage identities of all neurotoxin gene sequences detected in 105 ancient DNA datasets.

**Supplementary Figure 3.**
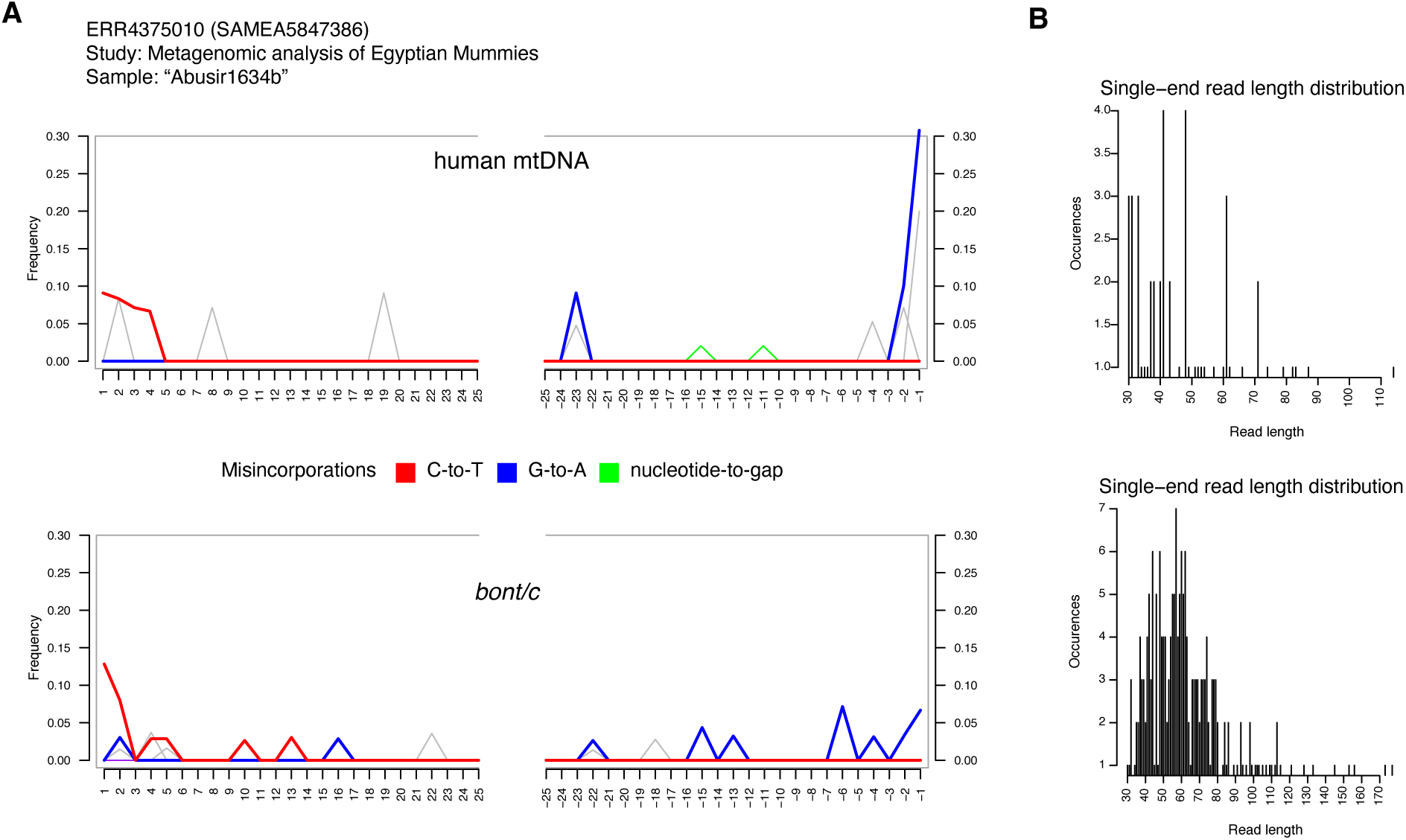
Additional MapDamage analysis of reads matching *bont/c* (bottom) and mitochondrial DNA (top) from an Egyptian mummy aDNA dataset. Reads were processed using leehom v1.2.18. (A) Misincorporation plots for human mtDNA (top) and *bont/c* (bottom) showing increases in C→T (and G→A) misincorporations at the edges of reads. (B) Distribution of read lengths mapping to human mtDNA (top) and *bont/c* (bottom). The plots were generated by trimming and merging reads with leeHom instead of fastp.

**Supplementary Table 1 –** dataset of toxin query sequences

**Supplementary Table 2 –** dataset of SRA datasets searched

**Supplementary Table 3 –** metadata associated with toxin-positive aDNA datasets

**Supplementary Table 4 –** query coverage of toxin gene matches in aDNA datasets

**Supplementary Table 5 –** percentage identities of toxin gene matches in aDNA data

